# Experimental Warming Alters Nitrogen Cycle in a Humid Tropical Forest

**DOI:** 10.1101/2024.12.10.627762

**Authors:** Parker M. Bartz, Iana F. Grullón-Penkova, Molly A. Cavaleri, Saima Shahid, Tana E. Wood, Benedicte Bachelot

**Affiliations:** Department of Plant Biology, Ecology and Evolution, Oklahoma State University, Stillwater, OK, USA; USDA Forest Service International Institute of Tropical Forestry, Rio Piedras, Puerto Rico; College of Forest Resources and Environmental Science, Michigan Technological University, Houghton, MI, USA; Plants, Photosynthesis and Soil, School of Biosciences, University of Sheffield, Sheffield, South Yorkshire, UK.

**Keywords:** nitrogen cycle, nitrogen fixation, microbial ecology, microbiome, plant biology, tropical ecology, global warming, hurricane, climate change, tropical forest

## Abstract

- Humid tropical forests typically contain high soil nitrogen (N), supporting rapid rates of primary productivity and recovery from disturbances. Nitrogen fixation is regulated by microbial communities, which can be free-living in the soil and leaf litter (asymbiotic) or in symbioses with plants.
- To investigate how warming affects asymbiotic and symbiotic components of the N cycle, we analyzed soil and leaf litter samples as well as annual seedling census data from an experimental warming field site in Puerto Rico.
- 16S and *nifH* sequencing revealed that warming significantly altered bacterial composition. Asymbiotic N fixation rates were 55% greater in soil (0.004 nmol N_2_/g/hr) and 525% greater (0.217 nmol N_2_/g/hr) in leaf litter from warmed compared to ambient plots. This increase in fixation was associated with changes in the N-fixing bacterial community. Under warming, N fixers experienced a 4.4-fold increase in growth rate compared to non-fixers, yet competition with neighboring N fixers eventually reduced N fixer growth. Seedling growth, especially of N fixers, initially increased following hurricane disturbances before declining.
- Together, our findings suggest that warming increases the flux of N from the atmosphere into this tropical forest, driving changes in both microbial and tree dynamics.

## Introduction

Tropical forests have an exceptional capacity to sequester carbon, support biodiversity, and cycle nutrients. Despite covering just 15% of Earth’s terrestrial surface, tropical forests account for two-thirds of global carbon (C) storage, one-third of global net primary productivity, and large portions of global nutrient cycling (Pan *et al*., 2013). These high rates of primary production and large biomass stocks, as well as rapid regeneration rates, are largely attributed to favorable climate, ample soil nitrogen (N), and quick recovery of the N cycle following disturbance (Brookshire *et al*., 2012). The interactions between N and C cycles within terrestrial ecosystems are likely to affect trajectories of atmospheric C concentrations and associated global climate changes (Thornton *et al*., 2009; Gerber *et al*., 2010; Zaehle and Dalmonech, 2011; Wårlind *et al*., 2014; Xu and Yuan, 2017). Thus, it is essential to study how N cycling in humid tropical forests will be impacted by climate change (Wood *et al*., 2012) and the altered regime of major disturbances such as hurricanes (Reed *et al*., 2020; Tunison *et al*., 2024).

Unlike other major plant nutrients, N is naturally brought into most ecosystems through the biological fixation of atmospheric N gas by microbes. N-fixing microbes are divided into two groups: asymbiotic (free-living) and symbiotic. Asymbiotic N-fixing microbes are hetero- or autotrophic bacteria, as well as some archaea, inhabiting water, soil, and leaf litter (Dynarski and Houlton 2018). Estimates from multiple tropical forests studies suggest that a substantial amount of N is fixed by asymbiotic microbes (e.g. Reed *et al*., 2007; Wurzburger *et al*., 2012; Matson *et al*., 2014): an estimated 0.459 – 0.714 kg N/ha/year in the humid tropical forest of the Amazon, for example (Moreira *et al*., 2021). Besides asymbiotic N-fixing microbes, others form a symbiotic relationship with specific plants and are housed in nodules in the roots of these plants. These N-fixing plants (N fixers) are relatively abundant in neotropical forests and provide a large potential source of N fixation (Taylor *et al*., 2017; Tamme *et al*., 2021). By bringing large inputs of N into these ecosystems, these N fixers may fuel rapid forest growth and carbon sequestration (Brookshire *et al*., 2019; Menge *et al*., 2019; Taylor *et al*., 2019).

At large scales, climate is considered a major driver of the microbial community function and composition (Castro *et al*., 2010; de Vries and Griffiths, 2018; Jansson and Hofmockel, 2020). Warming is especially important, as it can increase microbial processing, enzyme activity, turnover, and growth efficiency (Castro *et al*., 2010; Pajares and Bohannan, 2016; Wood *et al*., 2019). Warming conditions have long been thought to favor N fixation because it is a microbial process catalyzed by the enzyme nitrogenase (Houlton *et al*., 2007; Reed *et al*., 2011). Asymbiotic microbial N fixation rates have been found to increase in response to warming across many biomes, including tropical forests, although some studies reported only minor effects (Bai *et al*., 2013; Liu *et al*., 2016; Zheng *et al*., 2020). Extreme or chronic warming can inhibit asymbiotic N fixation, at least in high-latitude ecosystems (Rousk *et al*., 2018; Zheng *et al*., 2019), possibly by inducing water limitation of N-fixing microbes (Zheng *et al*., 2020). The long-term effects of warming on asymbiotic N fixation are still a critical point of study in lowland, humid tropical forests (Contosta *et al*., 2015; Pajares and Bohannan, 2016; Zheng *et al*., 2020).

In addition to influencing the functioning of microbes, warming may influence the composition of microbial communities. Warming can shift the asymbiotic microbial community to microbes adapted to higher temperatures and faster growth rates (Castro *et al*., 2010; Zhao *et al*., 2020) and can affect community composition via metabolic carbon or the soil nutrient pool (Ma *et al*., 2018). Experimental warming studies, which have largely been confined to temperate and higher-latitude ecosystems, have found that warming increased the richness of asymbiotic N-fixing microbial communities in the temperate grassland (Penton *et al*., 2015), and enhanced N-fixing microbial abundance and diversity in tundra ecosystem soils (Feng *et al*., 2019). While the relationship between asymbiotic N fixation rates and N-fixing microbial community composition has been investigated in terrestrial ecosystems, this relationship remains poorly investigated in tropical forests (Reed *et al*., 2010, Mirza *et al*., 2014, Pajares and Bohannan, 2016).

Warming may also affect the growth of N-fixing trees (Cusack *et al*., 2016; Nottingham *et al*., 2023). Much previous research on the effects of warming on plant growth in the tropics has centered on changes in photosynthesis and related physiology (Saxe *et al*., 2001; Cusack *et al*., 2016, Carter *et al*., 2021). Studies and meta-analyses have indicated that temperatures in tropical forests are likely already near optimal thresholds for plant metabolism and further warming may reduce photosynthetic rates by creating supraoptimal temperatures (Way and Oren, 2010; Yamori *et al*., 2014; Doughty *et al*., 2023). However, optimal temperatures for N fixation in plant symbioses can be higher than optimums for photosynthesis, suggesting plant C and N gain may be decoupled with respect to temperature (Bytnerowicz *et al*., 2022). Additionally, N fixation in trees may acclimate to growing temperatures in tropical forests (Bytnerowicz *et al*., 2022). Therefore, climate warming may increase the growth of N fixers in tropical systems.

N fixers affect the growth of neighboring plants and overall forest growth, but it there is limited data on how this relationship between N fixers and their neighbors will change with future warming. Decades of research has shown that N-fixing trees facilitate the growth of neighboring non-fixing trees (Uselman *et al*., 1999). It has been demonstrated that N can be transferred from tropical N-fixing trees to neighboring plants (Rao and Giller, 1993) via several pathways, including contributions to soil fertility by additions of organic N in plant litter, common mycorrhizal networks, and direct root-to-root transfer via exudation (Munroe and Isaac, 2014; Xu *et al*., 2019). However, N fixers can also inhibit growth of neighbors in secondary tropical forests (Taylor *et al*., 2017). These contrasting results might arise if N fixers facilitate growth of neighbors early in succession but inhibit growth later in succession, perhaps by taking light and other resources away from neighboring plants (Chapin *et al*., 2016; Taylor *et al*., 2017; Menge *et al*., 2019). These complex neighboring effects could be altered by climate change, especially if fixers and non-fixers respond differently to high temperature as previously discussed.

Finally, climate warming may affect how non-fixer and N fixer growth respond to additional disturbances, such as hurricanes. Hurricanes can directly increase tree damage and mortality (Walker 1995; Yap *et al*., 2016) and increase light availability to the less-damaged understory (Battaglia *et al*., 2001; Umaña *et al*., 2023). The understory also experiences a pulse of available soil nutrients, including N, associated with canopy-level defoliation (Lodge *et al*., 1991; Harmon *et al*., 1995; Xu *et al*., 2004; May and Oberbauer, 2021). As a result, N might become an excessive nutrient immediately following hurricane before rapidly dissipating (Hasselquist *et al*., 2010), changing how N fixers grow and interact with neighbors following hurricane disturbances. Studying how hurricane and warming influence plant demographics in N fixers and non-fixers is therefore important for determining future ecosystem resilience (Johnstone *et al*., 2016; Reed *et al*., 2020).

Given the contribution of tropical forests to the global N cycle, it is important to determine how N-fixing microbial communities and N-fixing trees in tropical forests will be affected by projected warming. In this work, we present the first comprehensive study of the effects of warming on N cycling dynamics in a tropical forest by utilizing a tropical field warming experiment to examine the two most important components bringing N into this system: asymbiotic N-fixing microbes and symbiotic N-fixing trees. To examine the effect of warming on asymbiotic microbes, we combined N fixation assay with DNA sequencing of soil and litter samples from the Tropical Responses to Altered Climate Experiment (TRACE), an *in-situ* field warming experiment in Puerto Rico. We hypothesized that warming would increase 1) N-fixing microbial community diversity and N fixation rates, indicating that warmed temperatures favor the growth and activity of N-fixing microbes, 2) growth of N-fixing seedlings compared to non-fixing seedlings, 3) the facilitative and competitive effect N fixers have on neighboring non-fixers and N fixers, respectively, and 4) the growth spike in N fixers following hurricane disturbances.

## Materials and methods

### Site description

This work took place at TRACE, in Puerto Rico. This study site is in a lowland subtropical wet forest located near the U.S. Forest Service Sabana Field Research Station, which is within the Luquillo Experimental Forest in northeastern Puerto Rico (18°18′N, 65°50′W). Located at 100 m elevation, the site has a mean annual temperature of 24°C with relatively low seasonal variation, and mean annual rainfall is 3,500 mm (Alonso-Rodriquez *et al*., 2022). The site contains clay-rich, acidic soils classified as Ultisols, and >60% of root biomass is concentrated in the top 10 cm soil layer. In the study area, the dominant canopy tree species are *Syzygium jambos*, *Ocotea leucoxylon*, and *Casearia arborea*, and the site is also dominated by the palm *Prestoea montana*. The most prevalent N-fixing tree species are *Inga laurina* and *Inga vera*. The TRACE experiment, which began in 2015, utilizes infrared (IR) heat to warm three 12 m^2^ plots within the forest to 4°C above ambient temperatures, which can be compared to three control ambient plots (Fig. 1; Kimball *et al*., 2018). Warming treatment started in September 2016 and continued until hurricanes Irma and Maria hit the island in September 2017, causing heavy damage to the site. Warming was halted for a year before restarting in September 2018. Prior to the hurricanes, surface soils (0-10 cm) in warmed plots were warmed 3.6°C above ambient soils (Kimball *et al*., 2018). After the hurricanes, the IR heaters were initially 2.5 m above the ground and moved to their max height of 4 m above the ground in July 2020. As understory vegetation height and foliage density increased, the warming effect in surface soils was reduced to ∼ +1°C (Tunison *et al*., 2024). These experimental conditions enable study the effects of warming and hurricane disturbance on various ecosystem processes in this tropical forest (Reed *et al*., 2020; Tunison *et al*., 2024).

**Fig. 1:**
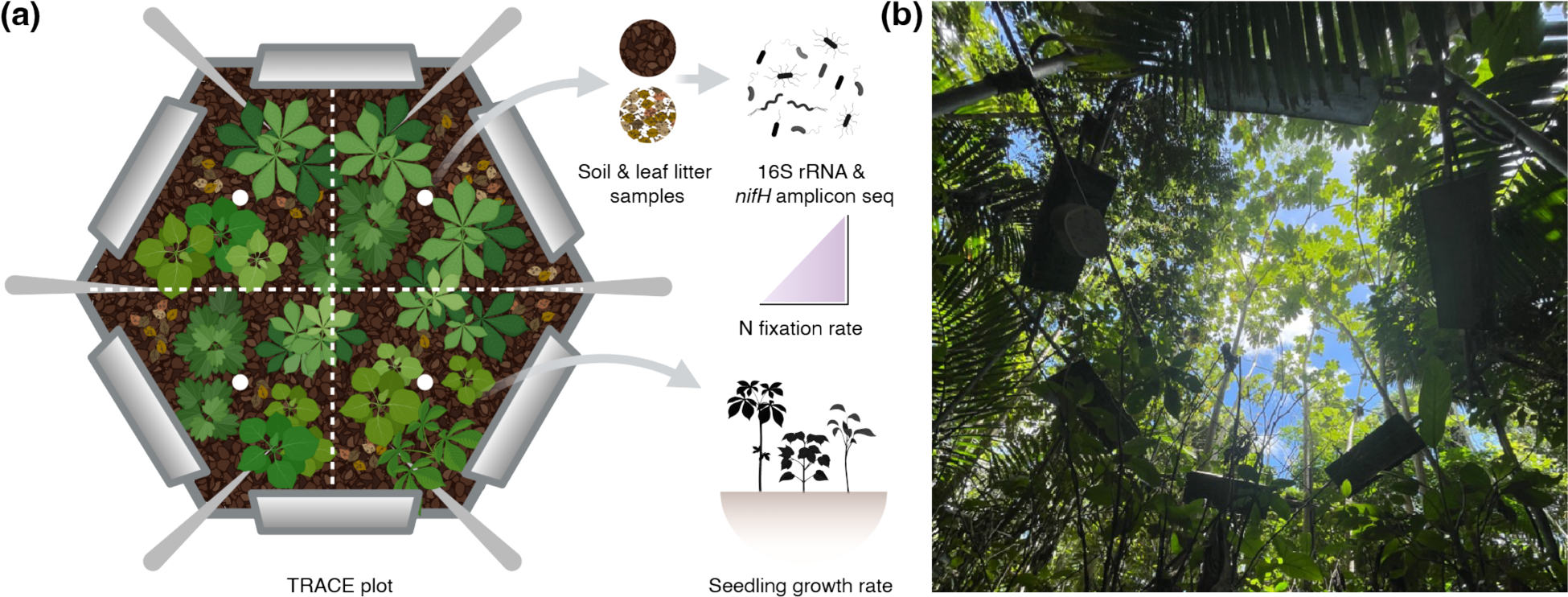
Experimental setup of the TRACE plots in the Luquillo Forest of Puerto Rico. The TRACE experiment utilizes infrared heaters to warm three 12 m^2^ plots within the forests. These warming plots are compared to three control plots experiencing ambient temperatures. (a) Diagram depicting top view of a TRACE plot used for experimental warming. Each TRACE plot is divided into four equal quadrants (represented by dashed lines). The circles at the center of each quadrant represent the approximate location of sites where soil and leaf litter samples were collected for N-fixation measurements and microbial sampling. (b) View from underneath one of the TRACE plots in the Luquillo Experimental Forest.

### Field collections

In July 2023, one leaf litter sample and one soil sample were collected from each of the 4 quadrants in each of the 6 plots (Fig. 1A). Using sterilized nitrile gloves, approximately 4 grams of leaf litter and 13 grams of soil were collected from the center of each quadrant. Four additional leaf litter and four additional soil samples were collected approximately 2 meters in a random direction away from two of the warmed plots and away from two of the control plots (for baseline ethylene concentration measurements). Each sample was immediately inserted into a 55-ml clear acrylic tube and returned to the on-site laboratory at Sabana Field Station, where an acetylene reduction assay was used to estimate asymbiotic N fixation rates (see N fixation rate section). Following this, samples were subsampled, and a portion was dried to calculate moisture content; these subsamples were then ground to calculate nitrogen, carbon, and phosphorous content (see C, N, and P content section). The remaining samples were stored in individual bags in a -80°C freezer before being shipped to Oklahoma State University for characterization of the microbial community composition (see microbial community composition section).

### N fixation rate

The fixation rates of 24 leaf litter and 24 soil samples were analyzed using an acetylene reduction assay (Hardy *et al*., 1968; Reed *et al*., 2010). Each 55-ml tube was sealed with a lid fitted with a septum, injected with 3.0 mL acetylene (creating approximately 10% headspace concentration), and vented to the atmosphere. The additional samples collected away from four of the plots, 8 in total, were not injected with acetylene so measurements of ethylene produced in the absence of acetylene could be determined. Eight blank samples were created by injecting acetylene into tubes without sample inside so measurements of acetylene reduced to ethylene in the absence of sample could be determined. After an approximately 18-hour incubation in the lab at ∼26°C, headspaces of all 64 samples (48 plot samples, 8 control samples, and 8 blank samples) were subsampled and injected into Exetainer tubes. These Exetainer headspace gas samples were transported to the USGS Southwest Biological Science Center (SBSC) in Moab, Utah, and analyzed for ethylene concentration on a gas chromatograph (Shimadzu GC-14A equipped with a Flame Ionization Detector, Shimadzu Corporation, Kyoto, Japan).

The baseline ethylene concentrations of the additional soil and leaf litter samples, and blank samples were averaged respectively. To account for ethylene production by soil (or leaf litter) alone, as well as for ethylene production in the absence of sample, the average baseline ethylene concentration of the additional soil (or leaf litter) samples and blank samples was subtracted from the baseline ethylene concentration of plot soil (or leaf litter) samples to obtain the adjusted ethylene concentration for each sample.

For plot soil samples, which were compact at the bottom of each tube, headspace volume was calculated via the measured height of empty space in each tube: from the soil surface to the bottom of the lid. For plot leaf litter samples, which were packed loosely, a headspace volume of 25 mL was assumed; this is 70% of the total volume for an average empty tube. Dry weights for each sample were calculated by dividing each sample’s total wet weight by their moisture content (calculated from weighing wet and dried subsamples) + 1. Incubation times for each sample were calculated by measuring time at acetylene injection and time at headspace sampling. The adjusted baseline ethylene concentrations were converted to parts per million (ppm) of ethylene using the calibration curve calculated from standard samples. Then, the nanomoles of ethylene produced per gram of sample per hour were calculated using the adjusted ethylene concentration (ppm), headspace volume (mL), dry weight (g), and incubation time (hr) of each sample.

Each mole of ethylene produced is equivalent to one mole of acetylene reduced. acetylene reduction rates were then converted to N fixation rates by dividing moles of ethylene produced by three (Reed *et al*., 2010).

### C, N, and P content

After the acetylene reduction treatment, a ∼2 g subsample was taken from each soil sample, and a ∼1 g subsample was taken from each leaf litter sample, and the remaining portion of each sample was frozen in a -80°C freezer. The subsamples were dried at 105°C for 48 hours. These subsamples were pooled together by type and plot, and then ground using a SPEX SamplePrep 8000D dual mixer/mill (SPEX SamplePrep LLC, Metuchen, NJ). All ground samples were sent to the USGS SBSC, and total C and N content was determined using an Elementar Vario Micro Cube (Elementar Americas Inc., Ronkonkoma, NY). The ground samples were then sent to the BYU Environmental Analytical lab, where they were subjected to nitric acid digestion in a Mars 6 Microwave (EPA method 3051A; CEM Corporation, Matthews, NC) and total phosphorus content was determined using a Thermo Scientific iCAP 7400 ICP-OES Analyzer (Thermo Fisher Scientific, Waltham, MA).

### Microbial community composition

Soil and leaf litter samples stored in the -80°C freezer were thawed in 4°C and homogenized. Genomic DNA was extracted from each of the samples in triplicate using a Qiagen DNeasy Power Soil Pro Kit (Qiagen, Hilden, Germany). Additionally, along with both the soil and leaf litter extractions, the DNA extraction kit was run on blank controls containing no sample. Extractions were pooled for each sample and stored at -80°C until downstream processing.

DNA extractions were used to characterize the full bacterial community using the 16S rRNA gene and the N-fixing bacterial community using the *nifH* gene. Amplification and sequencing were carried out at the University of Minnesota Genomics Center (UMGC). Specifically, 16S rRNA gene sequences were amplified by PCR using the 16S primer set 515F-806R, which amplifies the variable V4 region of the 16S rRNA gene. The *nifH* gene, which encodes the reductase subunit of nitrogenase, the enzyme that catalyzes the reaction which fixes N, has been widely used as a genetic marker to study the diversity and abundance of N-fixing microbes in gene sequencing studies (Pajares and Bohannan, 2016). *NifH* gene sequences were amplified in two separate PCR steps to minimize the formation of undesirable primer dimers. First, the *nifH* gene was amplified using the *nifH* primer set Ueda19F-R6 (Sigma-Aldrich, St. Louis, MO), a broadly inclusive pair (Angel *et al*., 2018). This was followed by PCR amplification using Nextera-adapted primers suited for Illumina sequencing.

16S and *nifH* gene amplicons underwent v3 Illumina MiSeq 2x300bp paired end sequencing. Soil and leaf litter sequence data were processed separately using the Mothur MiSeq standard operating procedure (Schloss *et al*., 2009; Kozich *et al*., 2013). Briefly, paired-end reads were assembled for each amplicon and filtered to the expected amplicon length (291 base pairs (bp) for 16S and 394 bp for *nifH*). The trimmed 16S sequences were aligned to the reference sequence file Silva 132 database. The trimmed *nifH* sequences were aligned to the *nifH* sequence database described by Gaby and Buckley (2014). For both 16S rRNA and *nifH* genes, chimeras were removed using chimera.vsearch (Mothur) and the sequences were grouped into operational taxonomic units (OTUs) at 97% sequence similarity. OTUs with only one observation in a dataset were removed using remove.rare (Mothur). For 16S, taxonomy was classified in Mothur using version 19 of the Ribosomal Database Project (RDP) training set, publicly released in July 2023. For *nifH*, representative sequences from each OTU obtained using get.oturep (Mothur) were taxonomically classified by Basic Local Alignment Search Tool analysis against the NCBI GenBank database.

### Seedling survey

In June 2015, all seedlings of at least 10 cm height present in the 6 plots were tagged, identified to species, and measured for height from ground level and stem diameter at ground level. After a year of drought (2015), the census was repeated annually starting June 2016 until 2023 (with the exception of 2020). Using the census data from 2016-2023, we calculated seedling growth rates (growth rate = log(x_final_/x_initial_)/days between measurements, where x = height or stem diameter). By investigating both height and diameter growth rates, we can detect shifts in growth strategies such as conservative versus acquisitive (Reich, 2014; Zhao *et al*., 2016; Fagundes *et al*., 2022). We also measured crowding within quadrants due to fixers and non-fixers, by separating the effects of overall crowding (number of seedlings in the quadrant) from the effects of crowding from fixers calculated as the proportion of neighbors that are fixers (number of fixer seedlings in quadrant/total number of seedlings in quadrant). Using the date of the censuses, we calculated the number of days since last hurricane to investigate how hurricane disturbance affects seedling growth rates.

### Statistical analyses

#### Microbial composition and N fixation

Community composition analyses were conducted in the R programming environment using the *gjam* and *vegan* packages (Clark *et al*., 2017; Oksanen *et al*., 2022). Each of the four bacterial communities was analyzed separately: soil 16S, soil *nifH*, leaf litter 16S, and leaf litter *nifH*. Before analyses, all OTUs appearing in the soil or leaf litter blank controls were removed from the soil and leaf litter OTU composition, respectively. Homogeneity in dispersion of sequence relative abundances in OTUs from the warming and ambient groups was confirmed using *betadisper* with the Bray-Curtis distance metric. Distance-based redundancy analyses (dbRDA) were applied to the OTU composition data using *capscale*, again with sequence relative abundances and the Bray-Curtis distance metric. The abiotic (treatment, moisture, C:N ratio, and P content) and biotic (proportion of N-fixing seedlings, Shannon Index, and species richness in the given quadrant) independent variables in the dbRDA model were z-transformed. Permutation tests with 999 permutations determined which of these independent variables had significant effects on overall (16S) and N-fixing (*nifH*) bacterial composition.

Generalized joint attribute modeling (GJAM), with plot as a random effect, was used to analyze how the treatment, C:N ratio, and P content affected the bacterial communities. The GJAM analyses were conducted using Gibbs sampler (a Markov chain Monte Carlo technique, Geman and Geman 1984) with 2000 iterations and a burn-in period of 200. Using this approach, we identified OTUs associated with a significantly decreasing or increasing relative abundance in the warming treatment. We jointly analyzed N fixation rate of the samples and N fixing bacterial community to investigate how N fixation was directly affected by the predictor variables and indirectly via changes in bacterial composition.

#### Seedling demography

To investigate how seedling growth dynamics changed due to their N fixing status and the major disturbances experienced by the plots (warming and hurricane disturbance), the growth rate (*g_i,s,t_*) in both height and diameter of an individual seedling *i* from species *s* under the treatment *t* (ambient and warming) was modeled with the following equation: *g_i,s,t_* ∼ N(µ = *β_t_x_i_* + σ_species_ + (σ_plot_ /σ_seedling_), σ), where *x* represents a matrix of independent variables: treatment, N-fixation status (whether the seedling is capable of fixing N), number of seedlings in the given quadrant, the proportion of N-fixing seedlings in the given quadrant, days since hurricane disturbance, seedling’s initial height, as well as two-way interactions between N-fixation status and the other covariates (except seedling initial height), and three-way interactions among treatment, N-fixation status, and the other covariates (except seedling initial height). Count variables in the matrix (i.e. number of seedlings in a quadrant and days since hurricane) were log- and then z-transformed, and the independent proportion variable (proportion of N fixers in a quadrant) was z-transformed after taking the arcsine of the square root. Both of these linear mixed-effects models (height and diameter growth rate) included species (σ_species_), plot (σ_plot_), and individual seedling (σ_seedling_) random effects to account for repeated measurements.

## Results

### Microbial composition and N fixation

The 16S amplicon library for soil samples recovered 25,094 Operational Taxonomic Units (OTUs) from 1,883,812 rRNA sequences after quality trimming, removal of rare OTUs, and removal of OTUs that appeared in blank controls. Each soil 16S sample yielded an average of 78,492 quality sequences. There were 673 unique bacterial genera identified, which were assigned to 27 bacterial phyla. The phyla Planctomycetota and Pseudomonadota dominated both sample types, with an average relative abundance of 25.73% and 18.13% respectively. The 16S amplicon library for leaf litter samples recovered 34,857 OTUs from 3,416,750 quality rRNA reads and each leaf litter sample yielded an average of 142,364 quality sequences. There were 773 unique bacterial genera identified, which were assigned to 28 bacterial phyla. The phyla Pseudomonadota and Actinomycetota dominated both sample types, with an average relative abundance of 38.74% and 16.47% respectively.

The *nifH* amplicon library for soil samples recovered 4,495 OTUs from 47,158 quality sequences (this total excludes two ambient plot samples with too few quality reads that were excluded from all analyses), and each soil sample yielded an average of 2,143 quality sequences. Among OTUs that were classified (99.16%), here were 208 unique genera identified, which were assigned to 16 bacterial phyla. The phyla Pseudomonadota and Thermodesulfobacteriota dominated both sample types, with an average relative abundance of 71.87% and 10.55%, respectively, in ambient plot samples, and 66.86% and 15.62%, respectively, in warming plot samples. The *nifH* amplicon library for leaf litter samples recovered 8,811 OTUs from 167,914 quality sequences and each leaf litter sample yielded an average of 6,996 quality sequences. 98.68% of OTUs were assigned to classified, cultured bacteria using BLAST. 223 unique genera were identified, assigned to 17 bacterial phyla. Again, the phyla Pseudomonadota and Thermodesulfobacteriota dominated both sample types, with an average relative abundance of 71.89% and 12.81%, respectively.

As hypothesized, warming significantly altered both the overall and N-fixing bacterial community composition in soil and leaf litter (Table S1, Fig. 2a-b, Fig. 3a-b). Generalized joint attribute modeling of the overall bacterial community identified 16 OTUs in the soil and 44 OTUs in the leaf litter that were significantly associated with the ambient plots (Fig. 2c-f). These OTUs also show contrasting responses to edaphic factors (C:N ratio and phosphorus, Fig 4a-b). In the soil, OTUs associated with the warming plots tended to be negatively associated with C:N ratio and positively associated with phosphorus content while the opposite was true for OTUs associated with the ambient plots (Fig 4a). Additionally, 71 OTUs in the soil and 12 OTUs in the leaf litter were significantly associated with the warming plots (Fig. 2c-f). In the leaf litter, OTUs associated with ambient or warming plots responded similarly with C:N ratio and phosphorus (Fig 4b); OTUs associated with warming plots were positively associated with soil C:N and phosphorus, and OTUs associated with ambient plots were negatively associated with soil C:N and phosphorus. These OTUs were distributed across several genera and classes. Notably OTUs in the class Spartobacteria and in the genus *Pseudonocardia* were substantially more abundant in warming plots in soil and leaf samples, respectively.

**Fig. 2:**
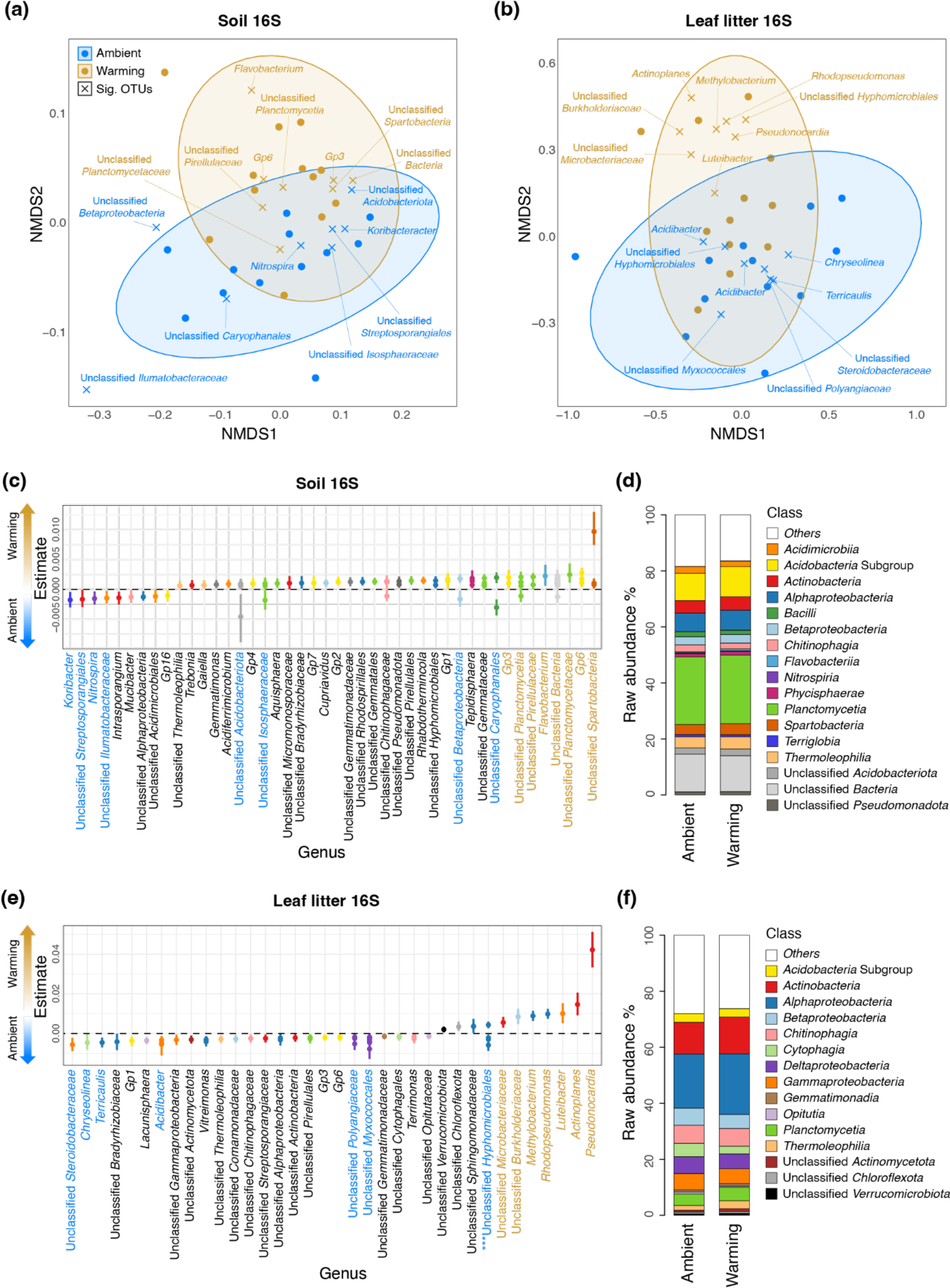
Differences in overall bacterial community composition elucidated by 16S rRNA gene sequencing of soil and leaf litter samples. NMDS plots depicting samples and significantly altered OTUs show that bacterial communities are distinct between ambient and warming treatments (a-b). Forest and raw abundance plots of significantly altered OTUs, listed by genus and colored by class, show how different genera respond to ambient versus warming conditions in soil (c-d) and leaf litter (e-f). The points on the x-axis represent the median of the standardized regression coefficients associated with warming treatment, derived from the posterior distribution of GJAM model parameters. Segments expand to the 95% credible intervals (CI). OTUs with 95% CI beneath 0 (dashed vertical line) are significantly more abundant under ambient conditions, and OTUs with 95% CI above 0 are more abundant under warming conditions. The 8 most highly altered OTUs in ambient and warming are highlighted. The asterisks indicate class where individual OTUs responded positively and negatively to the warming condition.

**Fig. 3:**
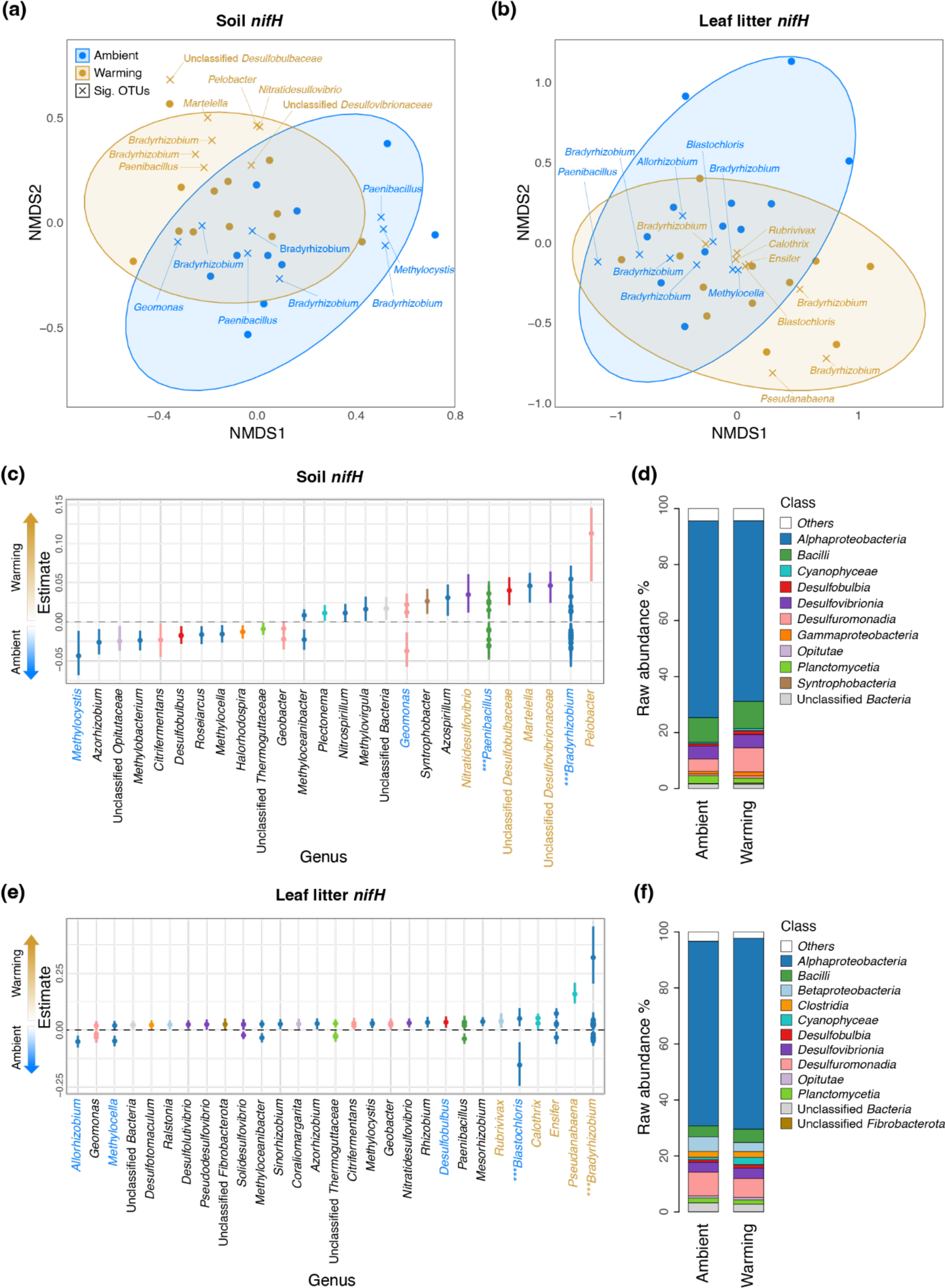
Differences in the N-fixing bacterial community composition elucidated by *nifH* gene sequencing of soil and leaf litter samples. NMDS plots show that bacterial communities are distinct between ambient and warming treatments (a-b). Forest and raw abundance plots of significantly altered OTUs, listed by genus and colored by class, show how different genera respond to ambient versus warming conditions in soil (c-d) and leaf litter (e-f). The points on the x-axis represents the median of the standardized regression coefficients associated with warming treatment, derived from the posterior distribution of GJAM model parameters. Segments expand to the 95% credible intervals (CI). OTUs with 95% CI beneath 0 (dashed vertical line) are significantly more abundant under ambient conditions, and OTUs with 95% CI above 0 are more abundant under warming conditions. The 8 most highly altered OTUs in ambient and warming are highlighted. The asterisks indicate class where individual OTUs responded positively and negatively to the warming condition.

**Fig. 4:**
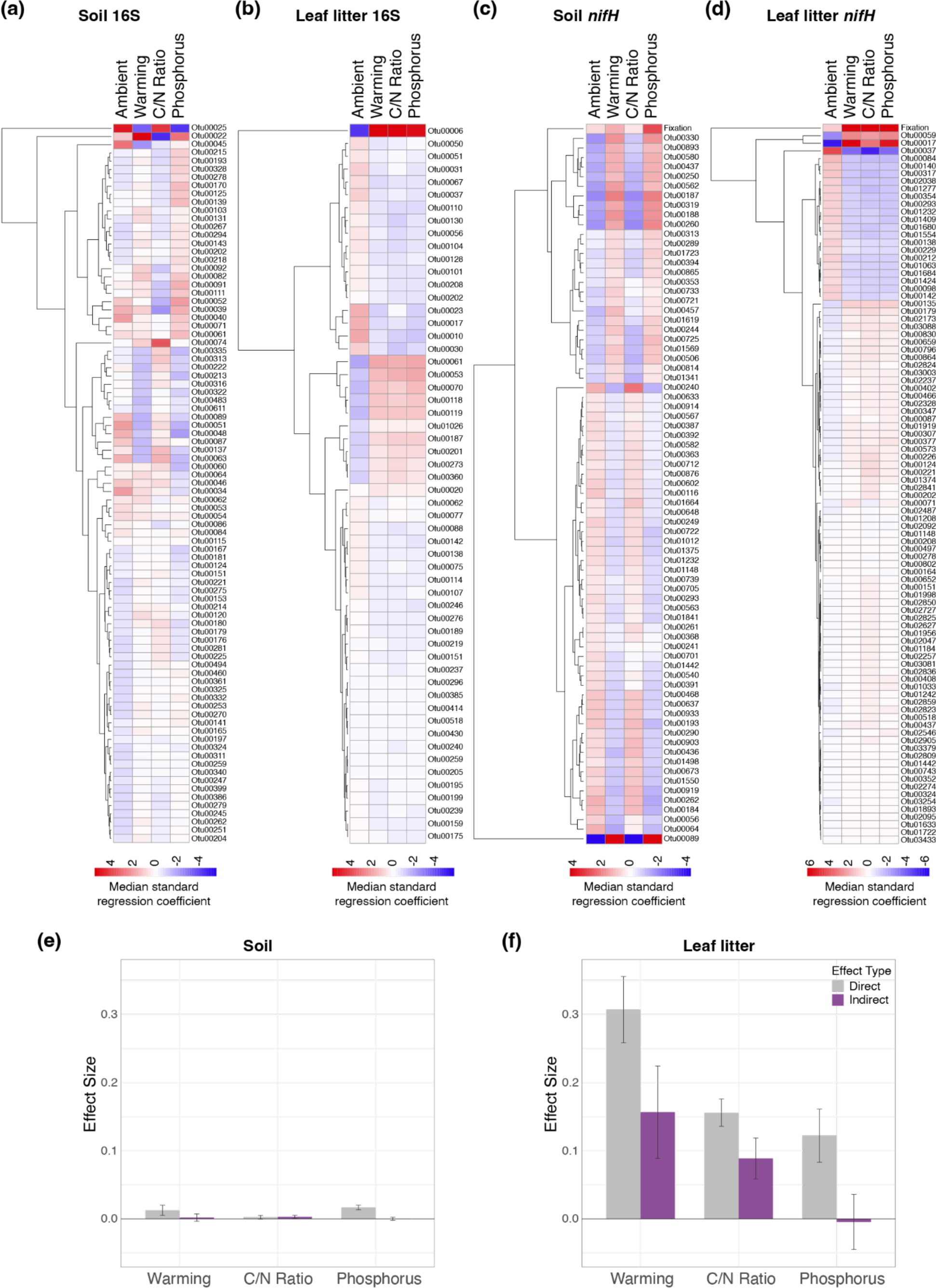
Individual and global responses to warming conditions as well as contribution of the bacterial community to N fixation. The heatmaps (a-d) represent the median of the standardized regression coefficients of the full GJAM models (warming treatment, C:N ratio, and phosphorus) for the OTUs that were the most affected by these variables for the overall soil and litter bacterial community (a-b) and the soil and leaf litter N-fixing bacterial community (c-d). The N-fixing bacterial community was modeled jointly with N fixation rates and therefore fixation is included in the response variables influence by the model in the heatmaps (c-d). Finally, the direct (gray) and indirect (purple) effects of warming, C:N ratio, and phosphorus content on N fixation rates in e) soil and f) leaf litter samples from the GJAM models. Bars reached the median effect and segments expand to the 95% CI.

Similarly, generalized joint attribute modeling identified many OTUs of N-fixing bacteria that were significantly associated with warming or ambient plots in the soil and leaf litter (Fig 3c-f, Fig. 4c-d). 47 OTUs in the soil and 20 OTUs in the leaf litter were significantly associated with the ambient plots, and 27 OTUs in the soil and 73 OTUs in the leaf litter were significantly associated with the warming plots. Notably, OTUs in the genera *Pelobacter* and *Pseudanabaena* were substantially more abundant in warming plots in soil and leaf samples, respectively. Several OTUs in the genus *Bradyrhizobium* were considerably more abundant in warming, but several were also more abundant in ambient. As with the overall bacterial community, OTUs associated with ambient or warming plots showed contrasting responses to soil C:N ratio and phosphorus (Fig 4c-d). In the soil, OTUs associated with the warming plots were positively associated with soil phosphorus and negatively associated with soil C:N ratio, and the opposite was found for ambient plot OTUs (Fig 4c). Finally, as with the overall bacterial community, OTUs in the leaf litter associated with the warming plots also responded positively to soil C:N ratio and phosphorus (Fig 4d).

In soil, N fixation rates increased an average of 0.004 nmol N_2_/g/hr, or 55%, under warming compared to ambient. The direct effect of warming on N fixation was significant, but the indirect effect, which is the effect of warming on N fixation mediated by change in N-fixing bacterial composition, was not (Fig. 4e). Thus, the overall effect of warming was not considered significant in the soil. In leaf litter, N fixation rates increased an average of 0.217 nmol N2/g/hr, or 525%, under warming compared to ambient. Both direct and indirect effects, and thus the overall effect, were significant (Fig. 4f). Specifically, N fixation rate was significantly increased by warming, and this effect was both due to the direct effect of warming on N fixation rate and the indirect effect of warming mediated by changes in the N-fixing bacterial community.

### Seedling demography

As hypothesized, fixers exhibited a significantly higher rate of height growth than the non-fixers in the warming plots (Fig. 5). Specifically, under warming there was a 4.4-fold increase in the rate of height growth in N fixers compared to non-fixers. However, this fixer growth advantage was only for height growth and in the warming plots. If warming was beneficial to fixer seedlings, it was generally harmful to non-fixers individuals (Fig. 5): the warming treatment marginally and significantly decreased non-fixer height and diameter growth, respectively.

**Fig. 5:**
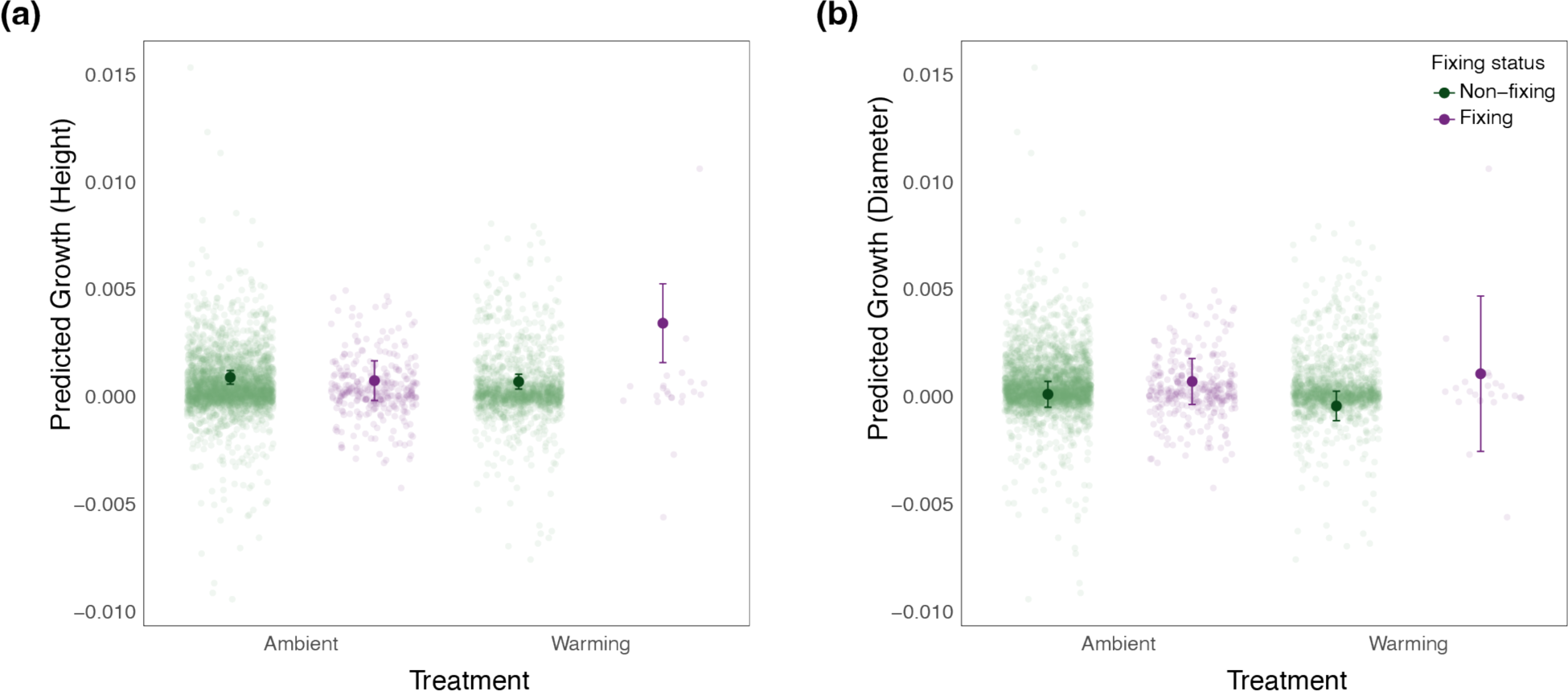
Growth model predictions in ambient and warming plots for non-fixing and N-fixing species. Both values of height (a) and diameter (b) growth predicted by the growth rate model are shown. Background dots represent individual, scaled seedling growth rates, and foreground dots represent predicted growth rates from the linear mixed-effects model. Segments expanding from the foreground dots represent the 95% CI.

As predicted, growth in height of non-fixer seedlings in ambient plots benefitted from a higher proportion of N fixers in the same quadrant (Fig. 6a). However, this effect was not present in warming. Also consistent with predictions, height growth of N fixers in warming plots decreased with a higher proportion of N fixer neighbors. Additionally, for N fixers in the warming plots, the decrease in height growth with increasing proportion of N fixer neighbors was significantly more drastic in seedlings with a greater total of neighboring seedlings. In contrast to height growth, N fixer diameter growth was not influenced by the proportion of nearby N fixers, in either the ambient or warming plots (Fig. 6b). Additionally, diameter growth of non-fixers in ambient plots decreased with increasing proportion of neighboring N fixers. Even so, a higher total number of neighboring seedlings was associated with less-decreased growth. Finally, non-fixers in warming experienced increased diameter growth with increasing proportion of N fixer neighbors, and this effect was significantly exaggerated by increasing total number of neighboring seedlings.

**Fig. 6:**
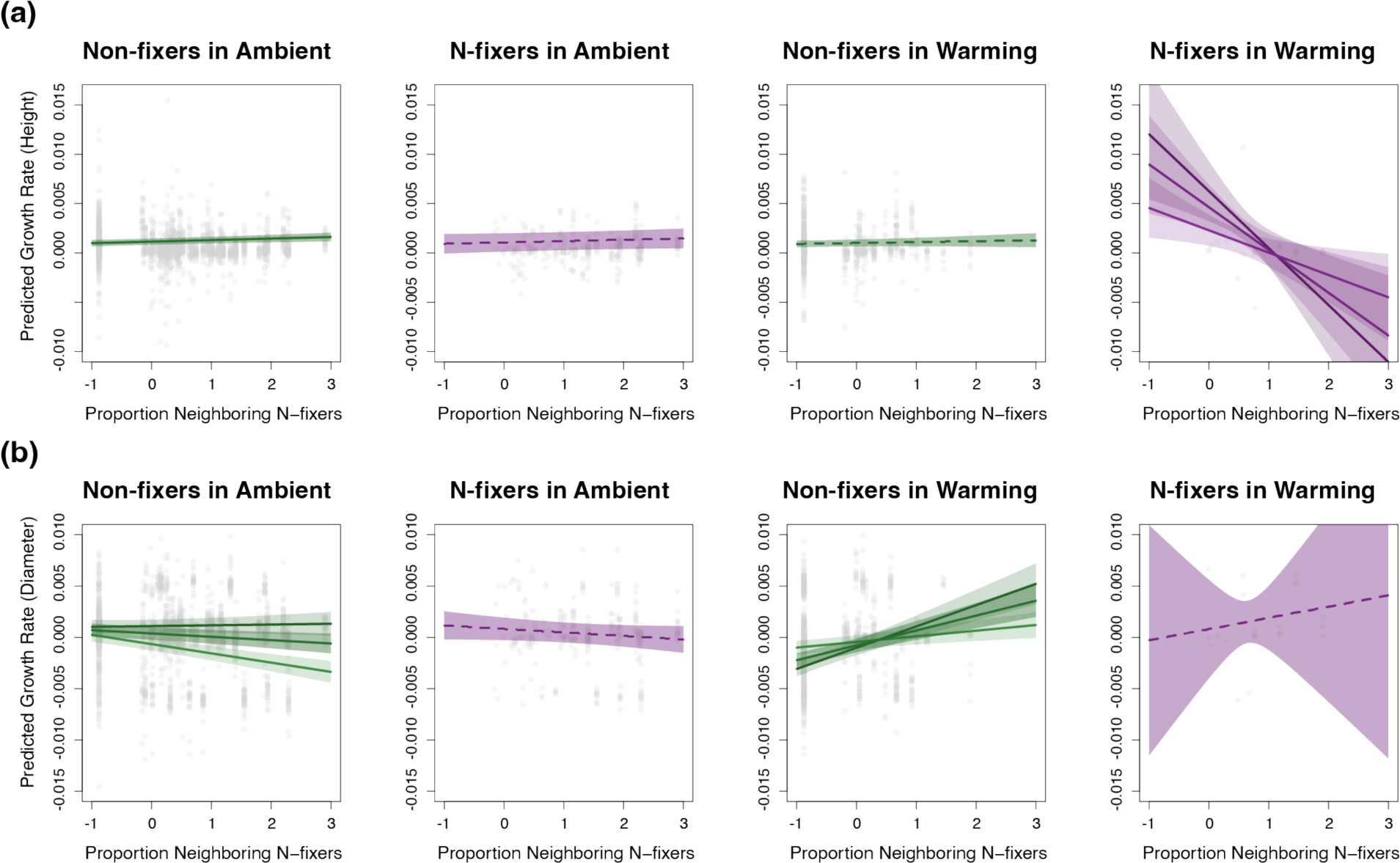
Values of height and diameter growth in ambient and warming plots predicted for non-fixing and N-fixing seedlings, as influenced by proportion of N-fixing seedlings in the quadrant and total number of seedlings in the quadrant. Background dots (gray) represent the scaled proportion of neighboring N fixers in the same quadrant as each seedling with a measured growth rate. Lines represent growth rates predicted for the proportion of neighboring N fixers by the linear mixed-effects model. Solid lines represent significant effects of neighboring N fixers on growth, while dashed lines represent non-significant effects. Panels with multiple lines indicate that the total number of seedlings in a quadrant also had a significant effect on growth. These lines represent growth rate predictions with low, medium, and high total number of neighboring seedlings, represented by light, medium, and dark coloring, respectively. Shading for all lines represents the 95% CI.

As hypothesized, as the number of days since hurricane increases, height growth of all seedlings significantly decreases, and this effect is significantly exaggerated in N fixers in ambient plots (Fig. 7a). Conversely, diameter growth increases with number of days since hurricane, although this effect is only significant in non-fixers (Fig. 7b).

**Fig. 7:**
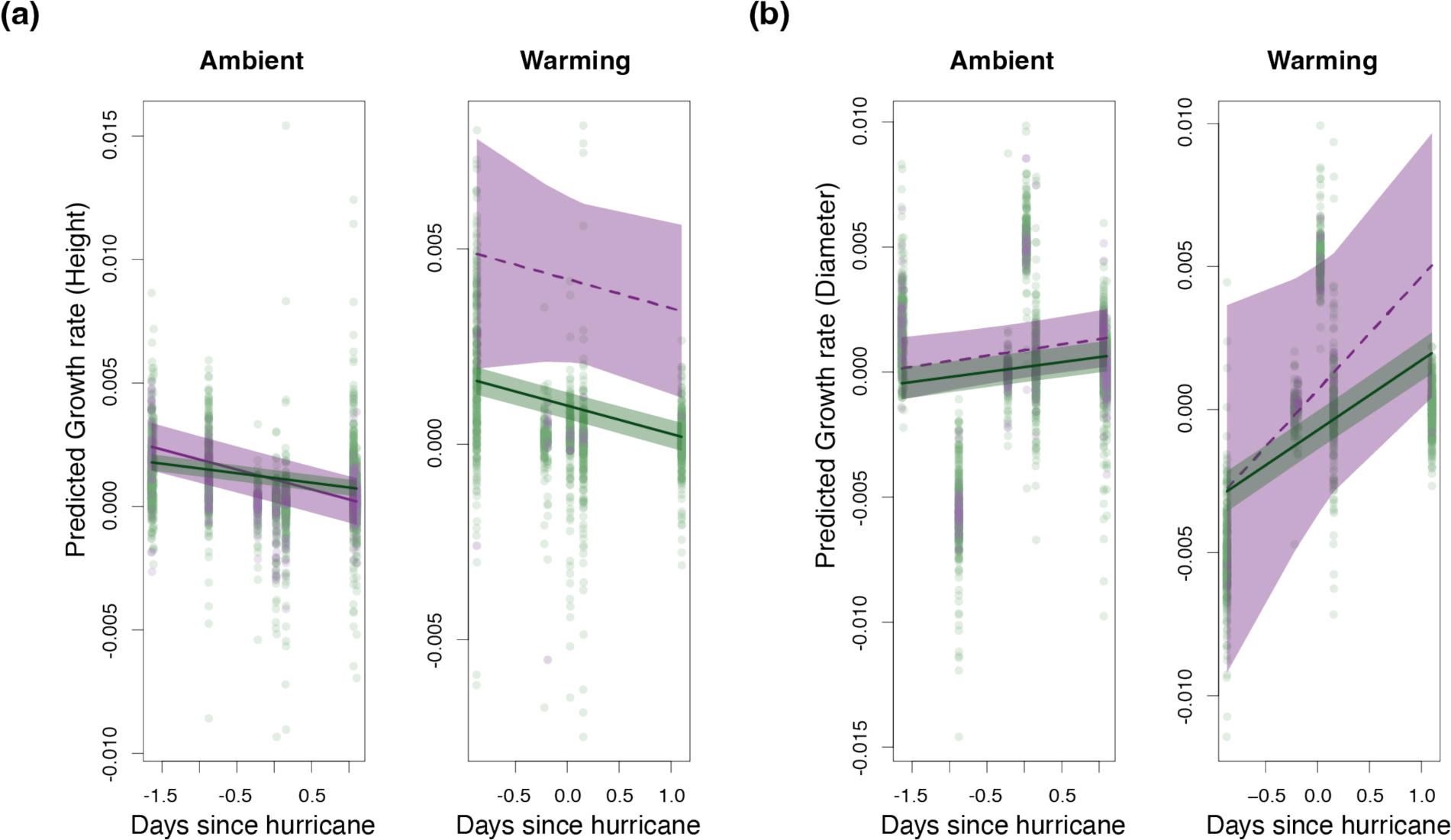
Values of height and diameter growth in ambient and warming plots predicted for non-fixing (green) and N-fixing (purple) seedlings, as related to number of days since hurricane disturbance, which is scaled. Background dots represent individual, scaled seedling growth rates. Lines represent growth rates predicted for the number of days since hurricane by the linear mixed-effects model. Solid lines represent significant effects of days since hurricane on growth, while dashed lines represent non-significant effects. Shading for all lines represents the 95% CI.

## Discussion

We provide the first comprehensive study on the effects of warming on asymbiotic and symbiotic N fixation aspects in a tropical forest by utilizing a tropical forest warming experiment, combining N-fixation assays, genetics, and long-term survey data. We found that warming increased asymbiotic N fixation activity and that this increase was partially mediated by change in the N-fixing bacterial community under warming. Both the overall and N-fixing bacterial communities were highly altered by warming, even more so than by other factors known to influence the asymbiotic bacterial community such as moisture, C:N ratio, and phosphorus. Enhanced asymbiotic N fixation activity under warming was echoed by our results in seedling growth, suggesting that symbiotic N fixation might also be enhanced. We found that N fixers had a growth advantage under warming, and crowding from neighboring plants nullified this advantage or even placed N fixers at a disadvantage. In both N fixers and non-fixers there was generally a shift from height investment to diameter investment following hurricane disturbance and from ongoing warming disturbance. Together, our results suggest that warming may increase N flux from the atmosphere to the ecosystem which if this N flux is coupled with a decrease in fertilizer use might help mitigating predicted increase in nitrous oxide emissions (Aryal *et al*., 2022).

### Warming significantly alters bacterial community composition and function

As hypothesized, warming increased N fixation activity: this increase is likely due to the effect of warming on both enzyme activity and bacterial composition. The direct effect of warming on N fixation was significantly positive in both the soil and leaf litter, and the effect of warming mediated by its effect on the N-fixing bacterial community was also significantly positive in the leaf litter. The differential responses in soil and leaf litter may be explained by previous findings that the two environments display different rates of N fixation, with leaf litter showing higher mass-based rates of N fixation (Reed *et al*., 2013; Van Langenhove *et al*., 2019) compared to soil. Therefore, leaf litter may have a high capacity for N fixation, allowing warming to increase fixation in leaf litter to a greater extent than in soil. The direct positive effect of warming on fixation observed in both environments may be due to increased enzyme activity and metabolism of N-fixing bacteria (Houlton *et al*., 2007; Reed *et al*., 2011; Wood *et al*., 2019), as the indirect effect accounts for the changing N-fixing bacterial community.

This indirect effect of warming is consistent with findings that temperature is a predominant parameter governing the N-fixing community (Zhao *et al*., 2020) and the N-fixing community is a strong predictor of N fixation activity (Berthrong *et al*., 2014). *NifH* gene composition doesn’t always correlate to fixation rates since bacteria containing *nifH* genes are not necessarily always fixing N (Feng *et al*., 2019), but in our study warming-induced shifts in N-fixing community composition did lead to increases in fixation. This is consistent with a previous study which found that variation in the N-fixing community accounted for differences in N fixation rates across a climate gradient which included tropical regions (Wu *et al*., 2020). Finally, while the activity and composition of N-fixing microbes in tropical soils have been found to be regulated by moisture, C:N ratio, and the availability of phosphorus (Pajares and Bohannan, 2016), our study shows that, even accounting for these parameters, warming generally had the strongest effects on activity and composition of N-fixing microbes in leaf litter.

The effects of warming on N fixation rates were in part explained by changes in bacterial composition. We found that artificial warming led to significant changes in both the overall and N-fixing bacterial community, as demonstrated by specific genera significantly associated with either the ambient or warming treatment. The taxa most highly associated with ambient plots in the soil include the nitrifying genus *Nitrospira* (Lücker *et al*., 2010) and the methanotrophic, N-fixing genus *Methylocystis* (Stein *et al*., 2011), which is also capable of nitrification. Additionally, Spartobacteria and *Pelobacter* were highly associated with warmed soils. Spartobacteria is an abundant, yet lesser-studied class of bacteria that, based on genome analysis, can likely degrade various complex carbohydrates but likely has no role in the N cycle (Herlemann *et al*., 2013; Cernava *et al*., 2017). *Pelobacter* is a lesser-studied N-fixing genus that is capable of producing sulfide (Li *et al*., 2021). The taxa most associated with ambient plots in the leaf litter include the order *Myxococcales,* which has predicted roles in carbon and potassium cycling (Dai *et al*., 2024), and the symbiotic N-fixing genus *Allorhizobium* (Egamberdieva *et al*., 2015). Taxa highly associated with warming in the leaf litter include *Pseudonocardia* and *Bradyrhizobium*. The genus *Pseudonocardia* is associated with certain plant and ant species and has been found to be capable of asymbiotic N fixation (Chen *et al*., 2019). *Bradyrhizobium* is a symbiotic N-fixing genus that is also capable of asymbiotic fixation and denitrification (Zhang *et al*., 2023). *Bradyrhizobium* may be warm adapted (Zhao *et al*., 2020), but some *Bradyrhizobium* OTUs were also associated with ambient plots, possibly because OTUs represent different bacterial strains.

The possibility of different strains or species of the same genus being affected differently by warming can be demonstrated by the example of Paenibacillus, which, in soil, has several OTUs associated with ambient plots and several associated with warming plots. The genus Paenibacillus is well-studied due to its ability to enhance plant growth by N fixation and P solubilization (Ryu *et al*., 2003). Further examination of specific OTUs in our study revealed that Paenibacillus species associated with ambient plots were different from species associated with warming plots. Warming may affect bacteria differentially even at the species level, emphasizing the need to further investigate in situ effects of warming on the bacterial community.

### Warming significantly alters N fixer demographics

Warming had strong effects on the growth dynamics of symbiotic N fixers. While warming 4°C above ambient temperatures was generally harmful to non-fixer seedling height and diameter growth, N fixer seedlings experienced increased growth. In contrast, Nottingham *et al*., (2023) found that seedling growth decreased in tropical soils heated 4°C, particularly in N fixer seedlings. This inconsistency may stem from different heating techniques used. Nottingham *et al*., (2023) utilized direct heating of soils in experimental plots, whereas in our study, plots were heated above soil level, with infrared heaters starting 1-2 m above soil level and rising to 4 m above soil level as understory vegetation height increased, resulting in the soils being heated to only 1°C above ambient temperatures (Tunison *et al*., 2024). Different warming temperatures can yield different growth responses. In a plant growth chamber study on the short-term effects of warming on two dominant tree species in a subtropical forest, Ghafoor *et al*., (2022) found that warming 2.3°C above ambient temperatures was beneficial for the height and diameter growth of the non-fixing species, while it was only beneficial for the diameter growth of the N-fixing species. Warming of 4.5°C above ambient temperatures reduced growth in both species, although it increased root nodule development in the N-fixing species.

Increased N fixer growth rates under warming suggest that warming affects symbiotic N fixation (Ghafoor *et al*., 2022; Nottingham *et al*., 2023), but evidence of symbiotic N-fixation rate changes is needed to show this conclusively. Although temperature effects root N exudation depending on the thermal optimums of species, climate warming has been considered to generally increase N exudation and play a role in sustaining increases in forest growth (Uselman *et al*., 1999). However, more recent results suggest that warming can also decrease root N exudation (Nottingham *et al*., 2023), and the relative importance of symbiotic N fixation in general has been questioned because it is facultative (Menge *et al*., 2009) and may decline to near zero in mature tropical rainforests (Barron *et al*., 2011; Van Langenhove *et al*., 2019). In future studies it will be important to measure the effects of warming on both N fixer growth and symbiotic fixation, as our knowledge of how temperature affects symbiotic N fixation, especially in the tropics, is primitive and based on imprecise methods that can be improved (Bytnerowicz *et al*., 2022). This will also be important when considering neighborhood effects in future warming studies on N fixer growth.

Our study also highlights differences in how N fixers and non-fixers respond to hurricane disturbances. Biogeochemical changes in tropical forest soils caused by hurricane disturbance will be shaped in concert with future warming (Reed *et al*., 2020). Our current study shows that, while N fixers likely receive a greater boost in growth after hurricane disturbance compared to non-fixers, the effect of warming, as well as the underlying mechanisms, of this growth response are still unclear. Over the longer term, studying the relationship between hurricane-induced changes in plant growth, symbiotic fixation, and warming will be important in determining future ecosystem resilience (Johnstone *et al*., 2016; Reed *et al*., 2020).

In both N fixers and non-fixers, there was a shift from height investment to diameter investment in the presence of other N fixers under warming, as plants potentially shift from acquisitive to conservative growth strategies. Acquisitive strategies take precedence when plants can maximize resource acquisition rates and focus on outcompeting others, leading to fast but vulnerable growth, especially height growth in the case of seedlings vying for sunlight (Zhao *et al*., 2016; Fagundes *et al*., 2022). Conservative strategies take precedence when limited resources or stressful conditions prioritize minimized resource loss, leading to slow but sturdy and safe growth, as in diameter growth (Reich, 2014; Fagundes *et al*., 2022). This suggests that resource competition and stress tolerance may be more critical factors under warming. Together, our analysis shows that, besides our hypotheses being supported to at least some extent, there is likely an additional trend of plants shifting from acquisitive to conservative growth following disturbance, including hurricane, and the plants in the warming plots are currently displaying more of a conservative growth pattern, having experienced consistent warming disturbance for over five years.

## Conclusion

Our study provides insight into the effects of warming on the asymbiotic N-fixing bacterial community, asymbiotic N fixation rates, and the growth of N-fixing symbioses in tropical forests, and therefore provides a uniquely integrated view of warming effects on N cycling dynamics in this biome. Our combined results suggest that warming will increase N flux from the atmosphere into the ecosystem while also increasing stress responses and competition in plants, possibly because warming elevates plant N demand even beyond microbial N production, resulting in conservative N utilization and growth in plants (Lie *et al*., 2021). Climate models predict that extremely high temperatures will occur more frequently in tropical regions than in other climatic regions (Anderson *et al*., 2012; Mora *et al*., 2013), so it is crucial to predict how warming-induced alterations in the N cycle in these regions will change ecosystem functions and contributions to global nutrient cycles in the future. Future studies should integrate the asymbiotic and symbiotic N fixation components explored in this study, in addition to symbiotic N fixation rates and other elements of the N cycle such as mineralization, nitrification, and denitrification, as well as N assimilation by plants and even N leaching.

## Acknowledgements

This study was funded by the US National Science Foundation (DEB-2120085, DEB-1754713) and the Department of Energy (DE-SC0012000, DE-SC-0011806, 89243018S-SC-000014, 89243018S-SC-000017, DE-SC-0018942, DE-SC0022095, 89243021S-SC-000076). We thank Tropical Responses to Altered Climate Experiment (TRACE) technicians Deyaneira Ortiz Iglesias, Laura Rubio Lebrón, Alberto Pastor Ibáñez, and William Mejía-García, as well as TRACE intern Katherine Pagán, for support in collecting, compiling, and organizing tree census data and processing samples collected from TRACE plots. Additional TRACE interns and volunteers also supported tree census work. We also thank USGS Southwest Biological Science Center biologists Cara Lauria, Shaina Lohman, and Kita Daly for support in collecting and processing soil and leaf litter samples from TRACE plots. We thank USGS biogeochemist Robin Reibold for support in determining sample nutrient contents and processing acetylene reduction assay samples. Finally, we thank Dr. Chelsea Murphy (Mathematical Sciences Department, Oklahoma State University) for support in using the Mothur program.

## Conflict of interest

The authors declare no conflicts of interest.

## Author contributions

PB and BB designed this current study. TEW, SCR, and MAC designed the warming experiment and run it. TEW, IFGP, BB oversee the annual seedling census, and the entire field season associated with this study. PB and SCR conducted the data collection for N fixation assays. SCR measured N fixation rates. PB analyzed all the data and wrote the first draft of the study. SS supported sequencing and data analysis. PB, BB, and SS generated figures for data visualization and edited manuscript. All authors provided regular feedback on the manuscript and data analyses.

## Data availability

All data and scripts associated with this study are available on request.

## Supplementary Information

**Table S1** The number of OTUs removed from each dataset because they appeared in the blank samples, the number of OTUs remaining for analysis, and the p-values of independent variables, determined by the distance-based redundancy analysis.

**Table S2** The fixed effects from the linear mixed-effect model analyzing the effects of various predictors on the rate of height growth.

**Table S3** The fixed effects from the linear mixed-effect model analyzing the effects of various predictors on the rate of diameter growth.

## Notes

### Competing Interest Statement

The authors have declared no competing interest.

### Summary of Updates

updated the PDF from the previous version to reflect new author list.

