## Supplementary material for "Experimental Warming Alters Nitrogen Cycle in a Humid Tropical Forest": Suplemental Tables

**Supplementary Information**

**Table S1:** The number of OTUs removed from each dataset because they appeared in the blank samples, the number of OTUs remaining for analysis, and the p-values of independent variables, determined by the distance-based redundancy analysis.

| | **Soil 16S** | **Leaf 16S** | **Soil *nifH*** | **Leaf *nifH*** |
|---|---|---|---|---|
| OTUs removed from analysis | 323 | 83 | 1063 | 1467 |
| OTUs kept for analysis | 25094 | 34857 | 4495 | 8811 |
| Treatment (effect on OTU composition) | 0.001 *** | 0.002 ** | 0.001 *** | 0.001 *** |
| Moisture | 0.009 ** | 0.032 * | 0.620 | 0.691 |
| C/N Ratio | 0.005 ** | 0.056 | 0.001 *** | 0.137 |
| Phosphorus | 0.015 * | 0.039 * | 0.005 ** | 0.567 |
| Proportion N fixers in quadrant | 0.945 | 0.060 | 0.090 | 0.423 |
| Shannon Index | 0.022 * | 0.107 | 0.019 * | 0.472 |
| Species Richness | 0.596 | 0.052 | 0.245 | 0.105 |

**Table S2:** The fixed effects from the linear mixed-effect model analyzing the effects of various predictors on the rate of height growth.

| Model Term | Estimate | Std. Error | T value | DF | P value |
|---|---|---|---|---|---|
| (Intercept) | 1.15E-03 | 1.62E-04 | 7.126387 | 24.2008 | 2.18E-07 |
| Warming | -1.63E-04 | 9.13E-05 | -1.78717 | 144.4955 | 7.60E-02 |
| N fixing | -6.01E-05 | 4.86E-04 | -0.1236 | 50.39075 | 9.02E-01 |
| Total seedlings in quadrant | -2.46E-05 | 3.24E-05 | -0.75921 | 1125.4 | 4.48E-01 |
| Proportion N fixers in quadrant | 1.67E-04 | 4.77E-05 | 3.504799 | 64.18369 | 8.39E-04 |
| Days since hurricane | -3.86E-04 | 2.48E-05 | -15.5332 | 4009.918 | 6.98E-53 |
| Initial Height | -7.79E-04 | 2.61E-05 | -29.7905 | 3261.01 | 1.07E-172 |
| Warming:N fixing | 3.30E-03 | 1.02E-03 | 3.24587 | 3553.221 | 1.18E-03 |
| Warming:Total seedlings in quadrant | 1.32E-04 | 5.07E-05 | 2.603872 | 2953.683 | 9.26E-03 |
| N fixing:Total seedlings in quadrant | -1.14E-04 | 1.97E-04 | -0.57986 | 3445.914 | 5.62E-01 |
| Warming:Proportio n N fixers in quadrant | -7.95E-05 | 9.59E-05 | -0.82901 | 406.4278 | 4.08E-01 |
| N fixing:Proportion N fixers in quadrant | -4.59E-05 | 1.06E-04 | -0.43385 | 1388.349 | 6.64E-01 |
| Total seedlings in quadrant:Proportion N fixers in quadrant | -5.66E-05 | 3.69E-05 | -1.53399 | 2890.901 | 1.25E-01 |
| Warming:Days since hurricane | -3.38E-04 | 5.57E-05 | -6.07658 | 4124.828 | 1.34E-09 |

| | | | | | |
|---|---|---|---|---|---|
| N fixing:Days since hurricane | -4.22E-04 | 7.74E-05 | -5.45672 | 3857.135 | 5.15E-08 |
| N fixing:Initial Height | -2.68E-04 | 9.44E-05 | -2.84172 | 3981.193 | 4.51E-03 |
| Warming:N fixing: Total seedlings in quadrant | 1.92E-03 | 5.92E-04 | 3.251297 | 4123.678 | 1.16E-03 |
| Warming:N fixing:Proportion N fixers in quadrant | -4.02E-03 | 1.37E-03 | -2.92631 | 4059.382 | 3.45E-03 |
| Warming:Total seedlings in quadrant:Proportion N fixers in quadrant | 8.20E-05 | 5.58E-05 | 1.469416 | 3441.536 | 1.42E-01 |
| N fixing:Total seedlings in quadrant:Proportion N fixers in quadrant | 1.28E-04 | 1.62E-04 | 0.790737 | 3714.812 | 4.29E-01 |
| Warming:N fixing:Days since hurricane | 4.02E-04 | 8.21E-04 | 0.489394 | 3385.813 | 6.25E-01 |
| Warming:N fixing:Total seedlings in quadrant:Proportion N fixers in quadrant | -1.85E-03 | 5.75E-04 | -3.2197 | 3718.549 | 1.29E-03 |

**Table S3**: The fixed effects from the linear mixed-effect model analyzing the effects of various predictors on the rate of diameter growth.

| Model Term | Estimate | Std. Error | t value | df | p value |
|---|---|---|---|---|---|
| (Intercept) | 2.00E-04 | 3.07E-04 | 0.651117 | 6.16053 | 5.38E-01 |
| Warming | -9.22E-04 | 2.18E-04 | -4.23383 | 611.0065 | 2.65E-05 |
| N fixing | 6.78E-04 | 4.96E-04 | 1.36753 | 101.67 | 1.74E-01 |
| Total seedlings in quadrant | 8.50E-04 | 7.67E-05 | 11.08379 | 2801.743 | 5.69E-28 |
| Proportion N fixers in quadrant | -4.25E-04 | 1.17E-04 | -3.6413 | 309.1631 | 3.18E-04 |
| Days since hurricane | 3.97E-04 | 5.74E-05 | 6.921784 | 4084.11 | 5.16E-12 |
| Initial Height | 3.76E-05 | 6.01E-05 | 0.626493 | 2066.791 | 5.31E-01 |
| Warming:N fixing | 7.25E-04 | 2.23E-03 | 0.324879 | 3277.034 | 7.45E-01 |
| Warming:Total seedlings in quadrant | -1.12E-03 | 1.18E-04 | -9.50128 | 3875.934 | 3.52E-21 |
| N fixing:Total seedlings in quadrant | -1.11E-03 | 4.37E-04 | -2.53525 | 3391.566 | 1.13E-02 |
| Warming:Proportio n N fixers in quadrant | 1.72E-03 | 2.25E-04 | 7.649909 | 1412.495 | 3.70E-14 |
| N fixing:Proportion N fixers in quadrant | -1.63E-04 | 2.37E-04 | -0.69078 | 3428.376 | 4.90E-01 |
| Total seedlings in quadrant:Proportion N fixers in quadrant | 4.73E-04 | 8.62E-05 | 5.485736 | 3944.261 | 4.38E-08 |
| Warming:Days since hurricane | 2.05E-03 | 1.27E-04 | 16.08563 | 3950.04 | 1.93E-56 |
| N fixing:Days since hurricane | 4.74E-05 | 1.79E-04 | 0.264364 | 4077.547 | 7.92E-01 |

| | | | | | |
|---|---|---|---|---|---|
| N fixing:Initial Height | -3.06E-04 | 2.18E-04 | -1.40849 | 3990.738 | 1.59E-01 |
| Warming:N fixing: Total seedlings in quadrant | 1.98E-03 | 1.36E-03 | 1.44845 | 4085.381 | 1.48E-01 |
| Warming:N fixing:Proportion N fixers in quadrant | 1.07E-05 | 3.17E-03 | 0.003384 | 4065.196 | 9.97E-01 |
| Warming:Total seedlings in quadrant:Proportion N fixers in quadrant | 2.64E-04 | 1.29E-04 | 2.042802 | 4060.407 | 4.11E-02 |
| N fixing:Total seedlings in quadrant:Proportion N fixers in quadrant | 7.02E-04 | 3.66E-04 | 1.918669 | 3914.242 | 5.51E-02 |
| Warming:N fixing:Days since hurricane | 1.46E-03 | 1.91E-03 | 0.767259 | 4063.788 | 4.43E-01 |
| Warming:N fixing:Total seedlings in quadrant:Proportion N fixers in quadrant | -1.67E-03 | 1.33E-03 | -1.25334 | 4075.563 | 2.10E-01 |
